# Crossmodal Pattern Discrimination in Humans and Robots: A Visuo-Tactile Case Study

**DOI:** 10.1101/775403

**Authors:** Focko L. Higgen, Philipp Ruppel, Michael Görner, Matthias Kerzel, Norman Hendrich, Jan Feldheim, Stefan Wermter, Jianwei Zhang, Christian Gerloff

## Abstract

The quality of crossmodal perception hinges on two factors: The accuracy of the independent unimodal perception and the ability to integrate information from different sensory systems. In humans, the ability for cognitively demanding crossmodal perception diminishes from young to old age.

To research to which degree impediments of these two abilities contribute to the age-related decline and to evaluate how this might apply to artificial systems, we replicate a medical study on visuo-tactile crossmodal pattern discrimination utilizing state-of-the-art tactile sensing technology and artificial neural networks. We explore the perception of each modality in isolation as well as the crossmodal integration.

We show that in an artificial system the integration of complex high-level unimodal features outperforms the comparison of independent unimodal classifications at low stimulus intensities where errors frequently occur. In comparison to humans, the artificial system outperforms older participants in the unimodal as well as the crossmodal condition. However, compared to younger participants, the artificial system performs worse at low stimulus intensities. Younger participants seem to employ more efficient crossmodal integration mechanisms than modelled in the proposed artificial neural networks.

Our work creates a bridge between neurological research and embodied artificial neurocognitive systems and demonstrates how collaborative research might help to derive hypotheses from the allied field. Our results indicate that empirically-derived neurocognitive models can inform the design of future neurocomputational architectures. For crossmodal processing, sensory integration on lower hierarchical levels, as suggested for efficient processing in the human brain, seems to improve the performance of artificial neural networks.

## 1 Introduction

Human behavior in the natural environment crucially depends on the continuous processing of simultaneous input to different sensory systems. Integration of these sensory streams creates meaningful percepts and allows for fast adaption to changes in our surrounding (1). The success of this crossmodal integration depends on two factors: The accuracy of the independent unimodal perception and the ability to integrate information from different sensory systems (2).

In a recent human behavioral study (Higgen et al. submitted; also posted at bioRxiv, doi: https://doi.org/10.1101/673491), we found that older participants show significant difficulties in a well-established visuo-tactile discrimination task compared to younger participants (3–5). This task combines the typical demands of crossmodal interactions. Participants have to detect simultaneously presented visual and tactile dot patterns and evaluate their congruency. With aging, performance decreases in several cognitive processes (6–9). The processing of unimodal sensory stimuli constitutes one major domain of this deterioration (10). However, our data revealed that difficulties of older participants go beyond a simple decline in unimodal stimulus detection. The data suggest that the integration of information from different sensory systems in higher-order neural networks might be one of the key reasons of poor performance of older participants.

Age-related alterations in human neural networks and their effects on local computing and long-range communication in the brain, which are needed for crossmodal integration, are not well understood (11–13). Causal assignment of altered neural function to behavioral changes is one of the great challenges in neuroscience. As the percentage of older people in the overall population increases, age-related declines gain more and more importance. Understanding the mechanisms of these declines is vital to develop adequate support approaches (see for example 13).

In the current study, we implemented a new approach by adapting our recent human behavioral study to an artificial neural network scenario. We employed embodied neurocognitive models to evaluate different hypotheses of the contribution of unimodal processing and crossmodal integration for the specific visuo-tactile discrimination task.

The design of high-performing artificial neural networks for crossmodal integration is likewise one of the most significant challenges in robotics. Therefore, the adaption of a human neurological experiment to an artificial scenario might help to establish common grounds in human and robotic research and the mutual exchange of theory. On the one hand, it will allow for an evaluation of the performance of artificial systems compared to humans with different abilities and help to develop more biologically plausible and performant artificial neural network (ANN) models. On the other hand, network models might help to understand the reasons for poor performance in older humans and can be a basis for the development of assistive devices.

## 2 Material and Methods

### 2.1 Visuo-tactile discrimination task in humans

In our human behavioral experiment, 20 younger (11 female, M = 24.05 years, SD = 2.50) and 20 healthy older volunteers (11 female, M = 72.14 years, SD = 4.48) performed an adapted version of a well-established visuo-tactile pattern discrimination task (3–5). In this task, participants had to compare Braille patterns presented tactilely to the right index fingertip with visual patterns presented on a computer screen (Figure 1). Patterns were presented synchronously, and participants had to decide whether they were congruent or incongruent.

**Figure 1.**
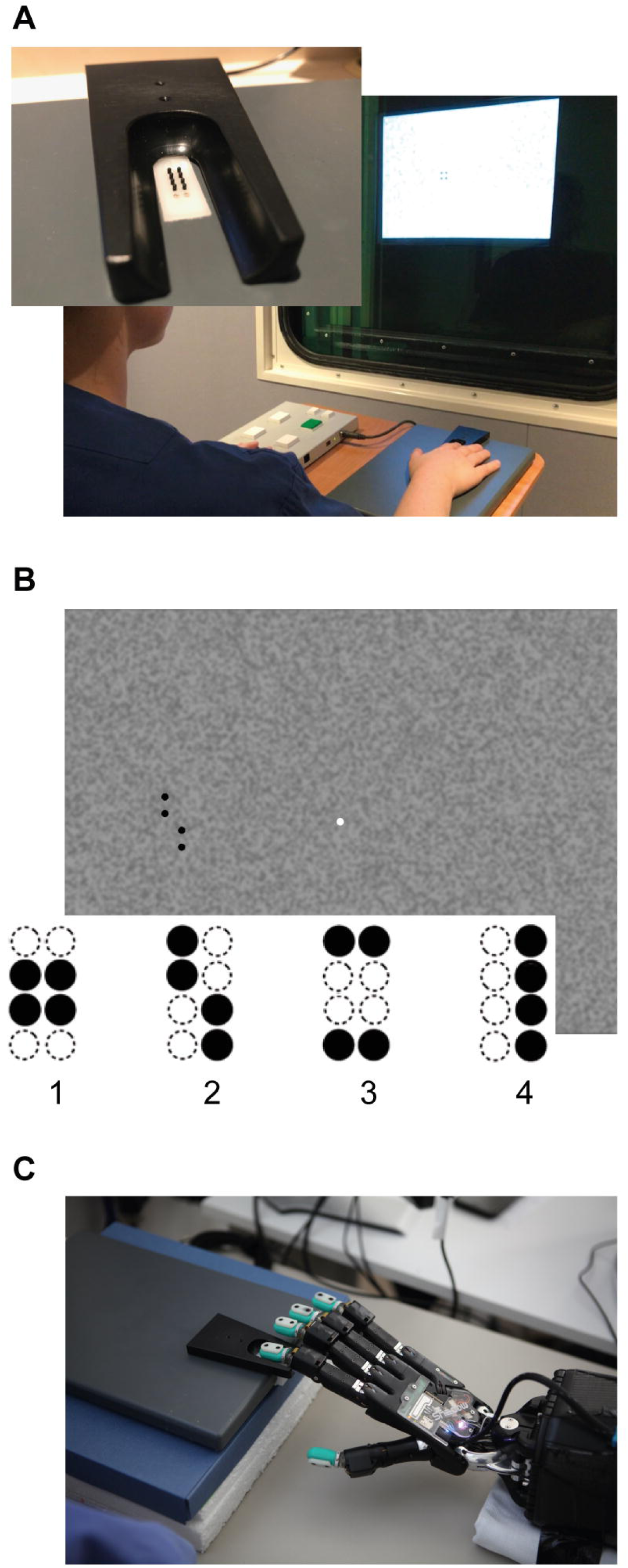
Experimental setup: **A:** Braille stimulator and setup of the human behavioral experiment. For tactile stimulation, the participants’ right hand was resting on a custom-made board containing the Braille stimulator (QuaeroSys Medical Devices, Schotten, Germany), with the fingertip of the right index finger placed above the stimulating unit. Visual patterns were presented on a monitor and the participants indicated whether both patterns were congruent or incongruent. The task was rendered more difficult by blending the visual pattern in the background noise and by reducing the actuated pin height. **B:** One example input for visual patterns with 100% intensity (i.e. full black). In both experiments, stimuli consisted of the same four patterns (1-4). **C:** Setup of robot experiment using a Shadow C6 Dexterous Hand. The BioTac tactile fingertip of the first finger of the hand is placed on the Braille stimulator.

Tactile stimulation was delivered via a Braille stimulator (QuaeroSys Medical Devices, Schotten, Germany, see Figure 1A), consisting of eight pins arranged in a four-by-two matrix, each 1mm in diameter with a spacing of 2.5mm. Each pin is controlled separately. Pins can be elevated (maximum amplitude 1.5mm) for a specific duration to form different patterns. Visual patterns were designed analogously to the Braille patterns and presented left of a central fixation point on a noisy background (Perlin noise; see Figure 1B) A set of four clearly distinct patterns was used in the study (see Figure 1B) to account for the diminished unimodal tactile perception of the older participants.

At the beginning of the experiment, unimodal stimulus intensities were adjusted individually based on an adaptive-staircase procedure with a target detection accuracy of approximately 80%. The adaptive-staircase procedure was performed in both modalities, to ensure comparable detection performance across modalities and between older and younger participants. Tactile stimulus intensity was adjusted by changing the height of the braille pattern (pin height). Visual stimulus intensity was adjusted by changing the patterns’ contrast against the background (gray level in % of black). Finally, participants performed the visuo-tactile discrimination task at the afore-defined unimodal thresholds.

The study was conducted in accordance with the Declaration of Helsinki and was approved by the local ethics committee of the Medical Association of Hamburg. All participants gave written informed consent. For a detailed description of the experiment, please see (Higgen et al. submitted; also posted at bioRxiv, doi: https://doi.org/10.1101/673491).

### 2.2 Robotic adaption

The setup described above was implemented in a robotic experiment (Figure 1C). For Braille stimulation, the same stimulator (QuaeroSys Medical Devices, Schotten, Germany, see Figure 1A) was used. The Braille stimuli were applied to the fingertips of a Shadow C6 Dexterous Hand (online: http://www.shadowrobot.com) equipped with BioTac tactile sensors (SynTouch LLC, BioTac Product Manual (V20), SynTouch LLC, California, Mar 2015. http://www.syntouchinc.com/wp-content/uploads/2017/01/BioTac\_Product\_Manual.pdf; (14)). The sensor surface of the BioTac closely matches the size and shape of a human finger, and it was possible to align and center the sensor onto the Braille stimulator without modifying the setup. To perceive the haptic stimuli of the Braille stimulator, the sensor can detect multiple contacts through indirect measurement. The turquoise rubber shell is filled with a conductive liquid and held in place around an inner rigid “bone”. When contacting an object, the rubber deforms, changing the overall pressure of the liquid (1 channel) and also the impedance between a set of electrodes patterned on the bone (19 channels). At the same time, the liquid temperature changes due to the contact (2 channels). Raw data from the sensor combines the measured pressure, temperature and impedances but it is difficult to interpret this raw data (16,17). Because the temperature conditions during recording remained stable, we omitted the respective sensor readings and fed the 20 other channels into an ANN to learn the mapping from raw data to applied Braille stimuli. As visual stimuli, we use the same visual stimuli employed in the human experiment. These stimuli are directly fed into the neural architecture without an intermediate sensor like a camera. As detailed below, the comparison with the human experiments relies on the exact gray values used in the stimuli; direct input of the images to the network avoids any level-shifts due to inconsistent camera exposure control. As the detection and classification of the tactile and visual stimuli require offline learning, the adaptive staircase procedure could not be used. Instead, we recorded patterns of different complexity in both modalities so that the required stimuli (corresponding to about 80% single-channel accuracy) could be presented to the trained ANNs after learning. We recorded several hours of raw sensor data from the robot, labeled with the presented tactile and visual patterns. In total, 3000 tactile samples were collected; matching visual stimuli are generated via image manipulation.

### 2.3 Computational models

To evaluate the influence of the actual crossmodal integration of high-level unimodal features in contrast to just comparing unimodal classification, we propose two neural architectures: The V-architecture (see Figure 2A) statically compares unimodal classification results. It consists of two separate networks that perform unimodal classification of the tactile and visual input pattern, respectively. Eventually, both classification results are compared in the final layer. In contrast, the Y-architecture (see Figure 2B) integrates high-level feature representations of both modalities. It also has two separate columns for unimodal feature extraction on the visual and tactile data. However, instead of performing a unimodal pattern classification, the extracted features are concatenated and further integrated by a series of dense layers, the stem of the Y-architecture. This network performs a late integration of crossmodal information. Empirical and automated optimization resulted in the following hyperparameters: For the visual columns, two convolution layers L1 and L2 after a batch-normalization step are followed by a pooling layer each (max-pool after L1, global max-pool after L2). For the V-architecture, the final dense layer of each arm uses soft-max activation to classify the four different outputs; in a second step these outputs are compared for equality. In the Y-architecture, the extracted high-level features are directly propagated. For the haptic modality, we use an MLP with three hidden layers (20 inputs from the BioTac sensor, three layers with 512 neurons each, followed by one softmax output layer with 4 neurons, corresponding to the 4 Braille patterns). Again, the last layer follows for the V-architecture only. Finally, the crossmodal integration in the Y-architecture is performed by a series of dense layers with a decreasing number of hidden units (64, 32, 16) followed by a binary softmax layer for same or different patterns.

**Figure 2.**
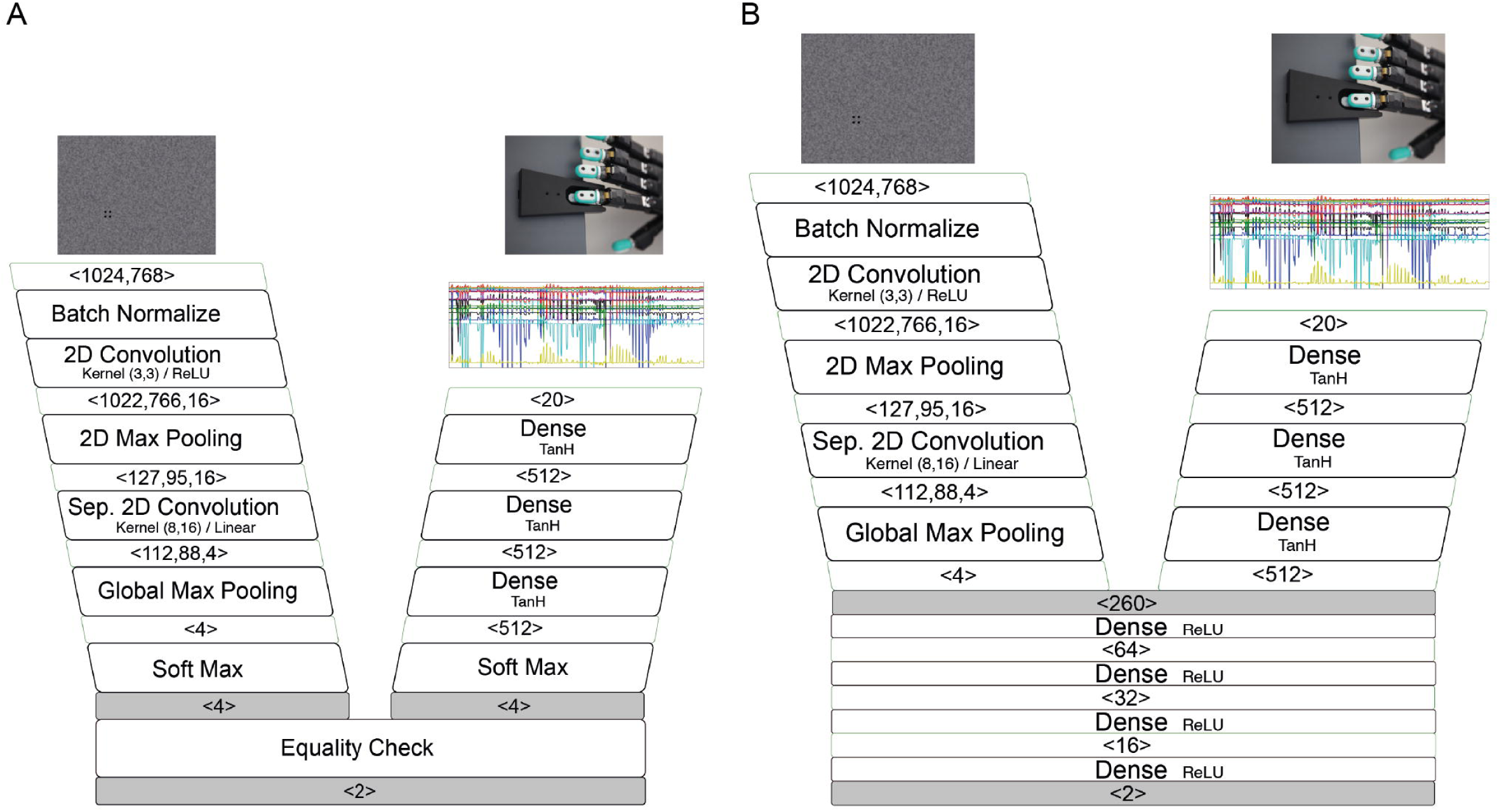
Structure of the neural architectures: **A:** Structure of the V-architecture. Visual (left column) and tactile data (right column) are processed separately and statically compared in the end. **B:** Structure of the Y-architecture. Both columns are first trained separately on visual and tactile data. Afterwards, a number of densely connected layers and a softmax output layer are added, and the network is trained again on the combined visual and tactile data.

### 2.4 Unimodal and crossmodal training

The training for both networks follows the same pattern: First, each unimodal column of the network is trained. In the case of the non-integrating V-architecture a static comparator follows. For the crossmodally integrating Y-architecture, a third training phase follows where the complete Y-architecture is trained. The tactile branch is pre-trained for 500 episodes and the visual network for 70 episodes. For the Y-architecture, at first only the integration network is trained for 70 episodes with frozen unimodal weights, and then the entire network is trained for another 70 episodes. For all training phases, the Adam optimizer with a learning rate of 0.001 and a batch size of 16 was used. The noisy visual input images are generated by placing one of four target Braille patterns (43×104px) randomly on one of 48 randomly generated background images (1024×768px, see Fig. 2 left). The background consists of a Perlin noise pattern with a gray range of between 40% and 60% of black (mean 53.7%). The stimulus intensity (i.e., gray level in % of black) of the pattern was selected to be between 47% and 100%. Samples were generated dynamically for each episode.

The Braille patterns become increasingly difficult to see for humans as the gray levels of the patterns blend with the gray levels of the background. On the robot, it might be possible to achieve even higher classification accuracy using classical computer vision algorithms and prior knowledge about how the data was generated. This would, however, undermine the goal to create controllable unimodal classification performances. Similar to the visual modality, a sufficient number of haptic samples with different pin heights were collected. Depending on the pin height and the unimodal network, different classification results can be achieved. Since only a limited amount of tactile training data could be collected on the real sensor, data augmentation was applied during training, generating each training sample by mixing two randomly selected tactile samples of the same pattern and pin height using a random interpolation/extrapolation factor between −50% and +150%. As in our human behavioral experiment, visual and tactile stimuli are paired so that the probability of both stimuli within a crossmodal sample pair representing the same symbol is 50%, equal to the probability of both stimuli representing different symbols. All test results for artificial neural networks were obtained through 10-fold cross-validation.

## 3 Results

### 3.1 Human behavior

In our human behavioral study, tactile and visual thresholds for a pattern detection accuracy of around 80% were estimated (See Table 1). Older participants showed higher thresholds for unimodal tactile and visual pattern detection compared to younger participants. In the crossmodal task older participants showed a significantly weaker performance compared to the unimodal condition when using the individual unimodal perception thresholds. In contrast, younger participants showed a stable performance of around 80% (See Table 1).

**Table 1.**
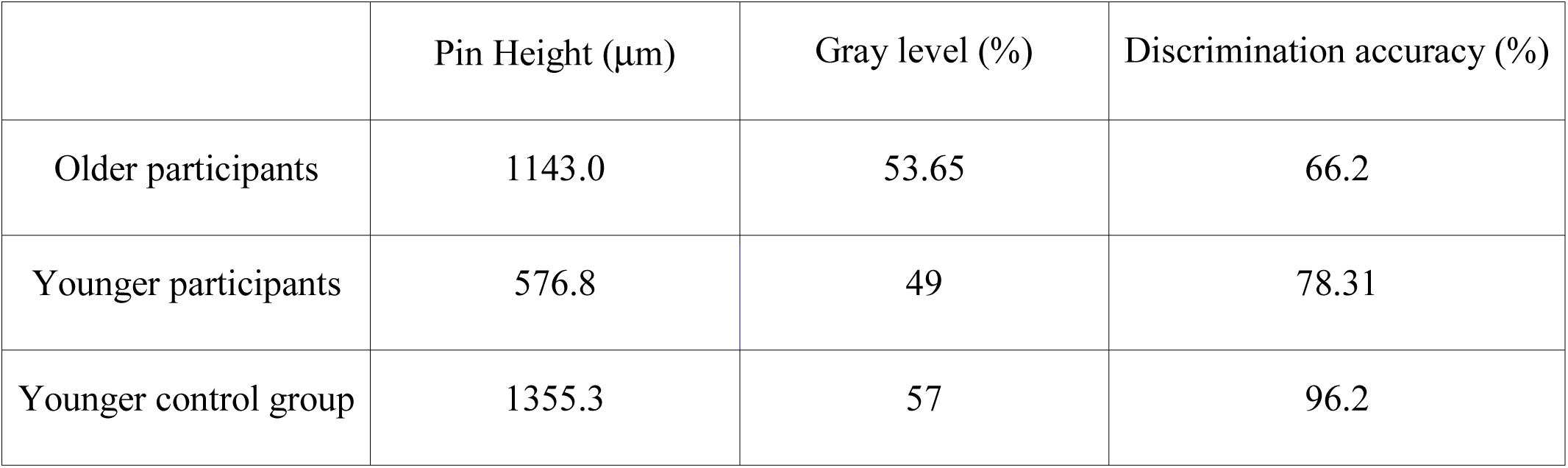
Performance of older and younger participants in the visuo-tactile discrimination task. Unimodal tactile (Braille pin height in μm) and visual threshold (gray level in % of black) for 80% detection accuracy, and performance on the visuo-tactile discrimination task at the unimodal thresholds, for younger and older human participants.

|  | Pin Height (μm) | Gray level (%) | Discrimination accuracy (%) |
| --- | --- | --- | --- |
| Older participants | 1143.0 | 53.65 | 66.2 |
| Younger participants | 576.8 | 49 | 78.31 |
| Younger control group | 1355.3 | 57 | 96.2 |

In a control experiment, younger participants showed a performance of 96.2% in the visuo-tactile discrimination task at thresholds comparable to the older group.

### 3.2 Artificial neural networks

To test the individual classification accuracy of both channels, and to compare them to the performance of the human participants in the original experiment, both models were fed inputs of varying difficulty (gray level for the visual channel, pin height for the haptic channel). A 10-fold cross-validation was performed by splitting the 3000 samples into 90% training and 10% test data. The results for the visual and tactile channel can be seen in Figure 3.

**Figure 3.**
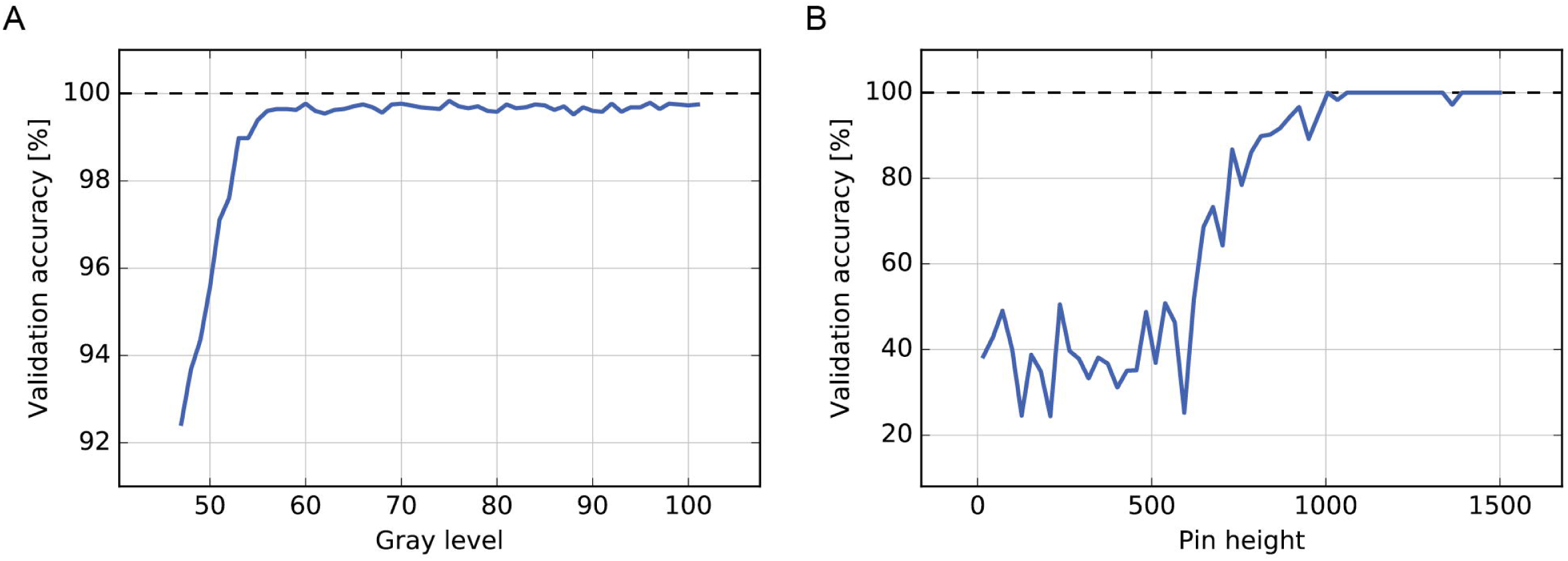
Unimodal performance of the artificial neural network: **A:** Unimodal performance of the visual pattern detection network which is also used as the visual branch in the V- and Y-architectures by gray level (% of black). **B:** Unimodal performance of the tactile pattern detection network which is also used as the tactile branch in the V- and Y-architectures by pin height (in μm).

In the visual condition, the classification accuracy was on average 99.04% and started dropping once the gray value of the pattern also appeared in the background image (values between 40% and 60%). Despite the increasing noise level, the performance of the network was high. At the lowest gray level (47 in % of black) pattern detection accuracy was at 91.42%. At a gray level of 54 (80% performance threshold of the older human participants) detection accuracy was at 98.85%. In the unimodal tactile condition, the artificial network showed a sigmoid learning curve, comparable to the human participants. A classification accuracy of 80% was reached around 730μm (82% accuracy).

The results for the multimodal V-architecture on the discrimination task are shown in Figure 4A. As expected, the performance of the network degrades when the channels are too noisy, but accuracy improves quickly as the signal quality (gray level, pin height) becomes better. The corresponding results for the Y-architecture that performs late crossmodal integration are shown in Figure 4B. For low stimulus intensities, the Y-architecture performs better than the V-architecture.

**Figure 4:**
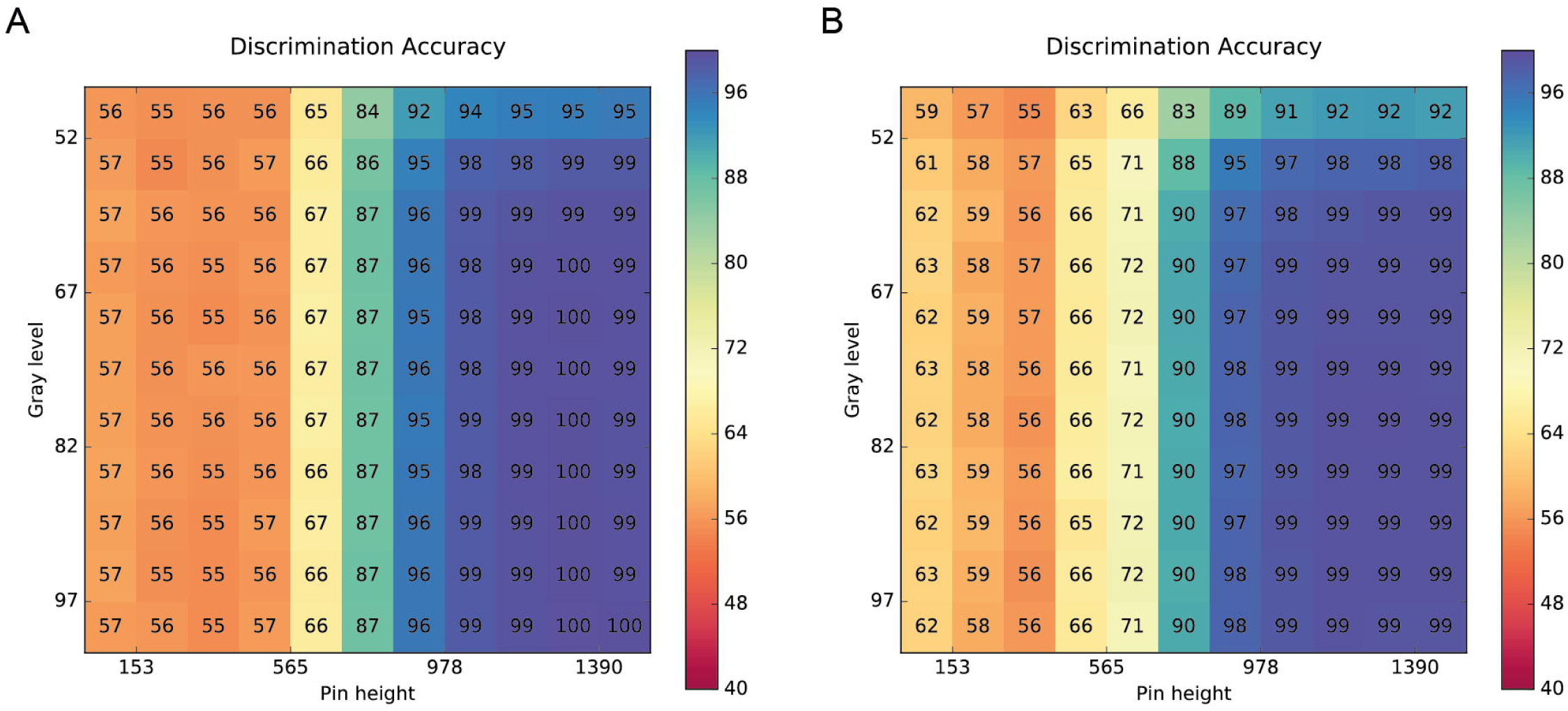
Crossmodal performance of the neural architectures: **A:** Performance of the V-architecture. Discrimination accuracy of the V-architecture (in %) by pin height and gray level. Parameters are the gray level (% of black) of the visual pattern and the active pin height (μm) of the Braille stimulator. **B:** Performance of the Y-architecture. Discrimination accuracy of the Y-architecture (in %) by pin height and gray level. Parameters are the gray level (% of black) of the visual pattern and the active pin height (μm) of the Braille stimulator.

When comparing the performance of the artificial neural networks in the visuo-tactile discrimination task to the human participants at their unimodal thresholds, it becomes obvious that both, the V- and the Y-architecture, outperform the older participants (see Table 2). Both show a performance above 95%, while older participants show a discrimination accuracy of 66.2%. The performance of the younger participants (96.2%) seems to be comparable to the ANN at these stimulus intensities. However, at the low stimulus intensities that constitute the thresholds of the younger participants, the performance of both networks is weaker than that of the younger participants (78.31%). At these stimulus intensities, the Y-architecture (62.5%) still outperforms the V-architecture (56.4%).

**Table 2.** Performance of the artificial neural networks compared to human participants. Discrimination accuracy of the artificial networks at tactile (Braille pin height in μm) and visual (gray level in % of black) signal levels comparable to the thresholds determined in the human experiments.

| Pin height (μm) | Gray level (%) | V-architecture accuracy (%) | Y-architecture accuracy (%) |
| --- | --- | --- | --- |
| 1143 | 54 | 98.6 | 97.5 |
| 593 | 49 | 56.4 | 62.5 |
| 1363 | 57 | 99.6 | 99.3 |

## 4 Discussion

This study aimed to investigate the transfer of a human behavioral experiment to an artificial neural network scenario and to compare the performance of different embodied neurocognitive models to performance of younger and older humans.

We implemented two artificial neural network models to evaluate different hypotheses of the contribution of unimodal processing and crossmodal integration to the visuo-tactile discrimination task. The first artificial network (V-architecture) implements a model for the integration of fully-processed results of the unimodal sensory streams. In contrast, the second network (Y-architecture) implements a model with an emphasis on the integration of information during crossmodal processing, integrating complex higher-level features from the unimodal streams.

The data show that in an artificial system, the integration of complex high-level unimodal features outperforms the comparison of independent unimodal classifications at low stimulus intensities even though the unimodal processing columns were identical. In our corresponding human behavioral experiment younger participants showed a stable performance in the crossmodal task at the unimodal thresholds while older participants showed a significantly weaker performance (Higgen et al. submitted; also posted at bioRxiv, doi: https://doi.org/10.1101/673491). The results suggest impaired mechanisms of crossmodal integration in the aged brain. Intriguingly, both datasets indicate that not only detection of unimodal stimuli but also mechanisms of integration are crucial for performance in crossmodal integration. The data of our two corresponding experiments now allow for comparing performances of the artificial neural networks and the human participants.

In the unimodal visual condition, we could not reach a target detection accuracy of 80% in the ANN. The data show that visual pattern recognition in artificial systems can be performed at very high levels. However, comparing this performance to the human participants is difficult, as stimuli were directly fed into the neural architecture without an intermediate sensor like a camera. Real-world data collection with adaptive camera exposure might lead to a weaker performance of the ANN in visual pattern detection. In the unimodal tactile condition, the performance of the ANN and the younger human participants was comparable, with a slight lead of the younger human participants. As the setup in the unimodal tactile condition was exactly the same in both experiments, the results show that state-of-the-art tactile sensors used can perform at a level almost comparable to humans (18). In both conditions, the ANN performed distinctively better than older participants.

In the crossmodal discrimination task, both artificial neural networks show high performance at high visual and tactile stimulus intensities (see Table 2). Performance is distinctly better compared to older participants and seems to be slightly better compared to young participants (see Table 1). However, for low stimulus intensities, the performance of the artificial networks lies below that of the younger human participants, despite the comparable or even higher performance in unimodal pattern detection. At low stimulus intensities, crossmodal integration appears to be more efficient and noise-resistant in younger human participants compared to the artificial systems. Comparing the performance of the artificial neural networks at these low stimulus intensities, the Y-architecture is performing better than the V-architecture. The Y-architecture seems to be more efficient compared to the V-architecture in integrating crossmodal stimuli at low intensities.

Taken together, the results for the artificial neural networks as well as human participants emphasize the importance of the mechanisms of integration for successful crossmodal performance. Early integration of incompletely processed sensory information seems to be crucial for efficient processing in crossmodal integration (19–22).

One might argue, that the V-architecture does not seem to be suitable to depict processes in the human brain as a decline in crossmodal integration processes is accompanied with poorer performance, as shown for the older participants in the human behavioral study. In contrast, the Y-architecture represents a more biological plausible network, approaching the efficient crossmodal processing in the young human brain. Still, our results suggest a superior mechanism for crossmodal stimulus processing in the young human brain. Further research is needed to answer the question of how young brains successfully integrate crossmodal information and which of these mechanisms can be adapted in artificial systems. It has been suggested that efficient stimulus processing in the human brain depends on recurrent neural networks and sensory integration on even lower hierarchical levels (23). Developing such approaches in future work might, on the one hand, improve the performance of artificial devices, but on the other hand, also give insights into the question which disturbances of the system lead to suboptimal functioning in the aged brain.

## 5 Author Contributions

FH: study idea, study design, data acquisition, data analyses, interpretation, preparation of manuscript. PR: study idea, study design, data acquisition, data analyses, interpretation, preparation of manuscript. MG: study idea, study design, data acquisition, data analyses, interpretation, preparation of manuscript. MK: study idea, interpretation, preparation of manuscript. NH: study idea, study design, interpretation, preparation of manuscript. JF: study design, interpretation, revision of manuscript. SW: interpretation, revision of manuscript. JZ: interpretation, revision of manuscript. CG: study idea, interpretation, revision of manuscript.

## References

1. Calvert GA. Crossmodal processing in the human brain: insights from functional neuroimaging studies. Cereb Cortex. 2001 Dec;11(12):1110–23.

2. Calvert GA, Spence C, Stein BE. The Handbook of Multisensory Processing. 2004 [cited 2018 Dec 7]; Available from: https://researchportal.bath.ac.uk/en/publications/the-handbook-of-multisensory-processing

3. Wang P, Göschl F, Friese U, König P, Engel AK. Long-range functional coupling predicts performance: Oscillatory EEG networks in multisensory processing. Neuroimage. 2019 Apr 5;196:114–25.

4. Göschl F, Friese U, Daume J, König P, Engel AK. Oscillatory signatures of crossmodal congruence effects: An EEG investigation employing a visuotactile pattern matching paradigm. Neuroimage. 2015 Aug 1;116:177–86.

5. Hummel FC, Gerloff C. Interregional long-range and short-range synchrony: a basis for complex sensorimotor processing. Prog Brain Res. 2006;159:223–36.

6. Heise K-F, Zimerman M, Hoppe J, Gerloff C, Wegscheider K, Hummel FC. The aging motor system as a model for plastic changes of GABA-mediated intracortical inhibition and their behavioral relevance. J Neurosci. 2013 May 22;33(21):9039–49.

7. Anguera JA, Gazzaley A. Dissociation of motor and sensory inhibition processes in normal aging. Clin Neurophysiol. 2012 Apr;123(4):730–40.

8. Guerreiro MJS, Anguera JA, Mishra J, Van Gerven PWM, Gazzaley A. Age-equivalent top-down modulation during cross-modal selective attention. J Cogn Neurosci. 2014 Dec;26(12):2827–39.

9. Gazzaley A, Cooney JW, McEvoy K, Knight RT, D’Esposito M. Top-down enhancement and suppression of the magnitude and speed of neural activity. J Cogn Neurosci. 2005 Mar;17(3):507–17.

10. Freiherr J, Lundström JN, Habel U, Reetz K. Multisensory integration mechanisms during aging. Front Hum Neurosci. 2013;7:863.

11. Hong SL, Rebec GV. A new perspective on behavioral inconsistency and neural noise in aging: compensatory speeding of neural communication. Front Aging Neurosci [Internet]. 2012 Sep 25 [cited 2016 Jul 18];4. Available from: http://www.ncbi.nlm.nih.gov/pmc/articles/PMC3457006/

12. Quandt F, Bönstrup M, Schulz R, Timmermann JE, Zimerman M, Nolte G, et al. Spectral Variability in the Aged Brain during Fine Motor Control. Front Aging Neurosci [Internet]. 2016 Dec 21 [cited 2018 Apr 18];8. Available from: https://www.ncbi.nlm.nih.gov/pmc/articles/PMC5175385/

13. Schulz R, Zimerman M, Timmermann JE, Wessel MJ, Gerloff C, Hummel FC. White matter integrity of motor connections related to training gains in healthy aging. Neurobiol Aging. 2014 Jun;35(6):1404–11.

14. Krawinkel LA, Engel AK, Hummel FC. Modulating pathological oscillations by rhythmic non-invasive brain stimulation-a therapeutic concept? Front Syst Neurosci. 2015;9:33.

15. Biomimetic tactile sensor for object identification and grasp control□: University of Southern California Dissertations and Theses [Internet]. [cited 2019 May 13]. Available from: http://digitallibrary.usc.edu/cdm/ref/collection/p15799coll127/id/475941

16. Lin C-H, Loeb GE. Estimating Point of Contact, Force and Torque in a Biomimetic.: 6.

17. Hsien Lin C, W. Erickson T, Fishel J, Wettels N, Loeb G. Signal Processing and Fabrication of a Biomimetic Tactile Sensor Array with Thermal, Force and Microvibration Modalities. In 2010. p. 129–34.

18. Dsouza R. The Art of Tactile Sensing: A State of Art Survey. International Journal of Sciences: Basic and Applied Research (IJSBAR). 2016 May 7;26:252–66.

19. Stein BE, Stanford TR. Multisensory integration: current issues from the perspective of the single neuron. Nat Rev Neurosci [Internet]. 2008 Apr [cited 2016 Sep 10];9(4):255–66. Available from: http://www.nature.com/nrn/journal/v9/n4/full/nrn2331.html

20. Molholm S, Ritter W, Murray MM, Javitt DC, Schroeder CE, Foxe JJ. Multisensory auditory-visual interactions during early sensory processing in humans: a high-density electrical mapping study. Brain Res Cogn Brain Res. 2002 Jun;14(1):115–28.

21. Kayser C, Petkov CI, Logothetis NK. Visual modulation of neurons in auditory cortex. Cereb Cortex. 2008 Jul;18(7):1560–74.

22. Kayser C, Logothetis NK. Do early sensory cortices integrate cross-modal information? Brain Struct Funct. 2007 Sep;212(2):121–32.

23. Ghazanfar AA, Schroeder CE. Is neocortex essentially multisensory? Trends Cogn Sci (Regul Ed). 2006 Jun;10(6):278–85.

